# PilD mutant is a new cheater in *Pseudomonas aeruginosa* quorum-sensing evolution

**DOI:** 10.1101/2022.11.22.517596

**Authors:** Huifang Qiu, Weijun Dai

## Abstract

Pathogen virulence is largely driven by the evolution and adaptation of the genome. LasR is a master regulator of quorum-sensing (QS) system. The QS-inactive LasR-null mutant regains QS activation in a laboratory evolution experiment through obtaining mutations in *mexT*, a gene encoding transcriptional regulator. How the QS-active LasR-MexT mutant continuously evolves is still unknown. Here we passaged the LasR-MexT mutant in casein broth and monitored its evolutionary trajectory. By examining the proteolytic phenotype of isolated single colonies, we found a protease-negative subpopulation arising from the parental population. The whole genome sequencing analysis revealed that these protease-negativ colonies bear mutations in *pilD* gene. In accordance with its role as a prepilin peptidase of type IV system, PilD mutants are defective in twitching motility. In conclusion, we identified a new PilD mutant cheater that evolved from the parental LasR-MexT mutant population. Our work demonstrates that population evolution is highly dynamics, with arising of distinct subpopulations containing different social behaviors.

## Introduction

*Pseudomonas aeruginosa* is an opportunistic pathogen that causes severe acute and chronic human infections (Gellatly and Hancock, 2013; Klockgether and Tümmler, 2017). A cohort of *P. aeruginosa* virulence factors is under the control of the quorum-sensing (QS) system (Papenfort and Bassler, 2016). QS is a bacterial cell-cell communication system that regulates the expression levels of hundreds of genes in a cell density-dependent manner. *P. aeruginosa* has two acyl-homoserine lactone (AHL) QS systems, Las and Rhl QS systems. LasR is a master regulator of Las QS system. In general, LasR-null mutant results in the inactivation of QS system.

Intriguingly, many LasR-null *P. aeruginosa* clinical isolates remain QS activity, producing QS molecules and QS-controlled virulence products, such as C4-HSL and pyocyanin (Bjarnsholt et al., 2010; Feltner et al., 2016). Through evolution experiments, mutations in *mexT* gene were found to be responsible for reverting QS activity in a laboratory strain PAO1 LasR-null mutant (Kostylev et al., 2019; Oshri et al., 2018). The *mexT* mutations down-regulates the expression of the MexEF-OprN efflux pump genes, which regulates the QS circuit (Köhler et al., 2001; Tian et al., 2009b).

To capture the evolutionary trajectory of the QS-reprogramming LasR-null revertant, we evolved a PsdR-LasR-MexT mutant strain in casein broth and monitor the social behavior of isolated single colonies. After one-month continuous passage, protease-negative clones appeared from the parental population. These clones bear mutations in *pilD*, a gene encoding the prepilin peptidase of type IV system. PilD mutants cheat cooperative parental population because they do not produce extracellular proteases. As PilD mutations disrupt the export of many virulence factors as well as the pilus assembly, the arising of PilD mutants expect to adversely affect the whole population virulence.

## Results

### Screen for protease-negative variants from PsdR-LasR-MexT mutant

Mutations in *mexT* are known to revert QS activity in the LasR-null mutant of *P. aeruginosa* strain PAO1 (Kostylev et al., 2019; Oshri et al., 2018). To further reveal the evolutionary trajectory of QS-active LasR-null revertants, we evolved the LasR-MexT mutant. As cells evolving in casein broth obtain *psdR* mutations for cellular dipeptide metabolism regulation (Asfahl et al., 2015), we initiated the evolution with a constructed triple mutant PsdR-LasR-MexT. Cells were evolved in casein broth, where production of extracellular proteases are necessary for cell growth. Two replicate mutant cells were passaged in casein broth every 5 day and screened for colonies deficient in protease-secretion. After 30-day evolution, protease-negative colonies appeared in one cell line, reaching to about 20% in the evolving population. These colonies exhibited LasR-null-like proteolytic phenotype, suggesting they obtained genomic mutations that impaired the extracellular protease production. These variants utilized the proteases from parent PsdR-LasR-MexT cells, and thus could be defined as cheaters.

### Identification of mutations responsible for protease-secretion inactivation

To identify mutations responsible for protease-secretion inactivation in those mutant variants, we next selected protease-negative clones for whole genome short read resequencing (WGS). The WGS analysis revealed short fragments of deletions in *pilD* (Table S1), a gene encoding a prepilin peptidase of Type 4 system. No other genomic mutations were identified in tested clones. We next amplified and sequenced fragments across the *pild* coding region in other protease-negative clones. These clones all shared mutations in *pilD*, suggesting mutated *pilD* resulting in protease-secretion inactivation.

### Phenotypic feature in PilD mutants

To characterize the phenotypic consequences of PilD mutants, we constructed the PsdR-LasR-MexT-PilD mutant. Consistent with the phenotypes of screened colonies, PsdR-LasR-MexT-PilD is protease-deficient (Fig. 1). On the other hand, As PilD is a prepilin peptidase responsible for removing the signal peptide of the prepilin pilA in the type IV system (Pepe and Lory, 1998), mutations in *pilD* were expected to paralyze the system. We thus sought out to examine their twitching motilities, a typical feature of type 4 system. As expected, those protease-deficient clones all lacked twitching motilities but not for the parental PsdR-LasR-MexT mutant. In conclusion, our results show that PilD mutant is novel cheater containing a disrupted type IV system.

**Figure 1.**
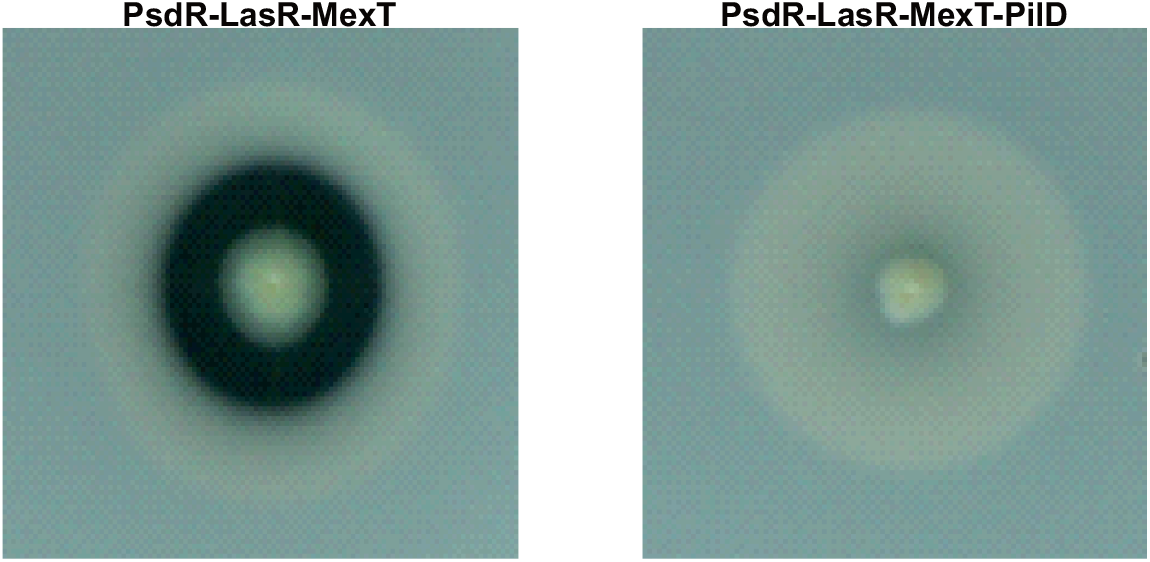
Protease activity of PilD mutants. Equal numbers of bacteria of indicated strains were spotted on skim milk plates to estimate proteolytic activity. Pictures were taken 24 h later. PsdR-LasR-MexT, triple mutant; PsdR-LasR-MexT-PilD, deletion of PilD from PsdR-LasR-MexT mutant.

## Discussion

In the present study, we characterized the evolutionary trajectory of a QS-active.PsdR-LasR-MexT mutant strain by continuous passage in casein broth. We found that mutant subpopulations arising from parental PsdR-LasR-MexT population contain mutations in *pilD* gene. Similar to the well-known typical cheater LasR mutant (Diggle et al., 2007; Sandoz et al., 2007), *pilD* mutants do not secret proteases and have to exploit extracellular proteases of neighboring cells. Unlike LasR as a master regulator of QS closely involving in QS circuit (Papenfort and Bassler, 2016), PilD encodes a prepilin peptidase of a type IV system and cleavages the signal peptide of the prepilin pilA prior to pilus assembly (Pepe and Lory, 1998). Therefore, further evolution of a QS-active population appears a novel non-QS cheater independent of QS regulatory pathway.

The arising of PilD mutant subpopulation may have impacts on the population virulence. PilD controls the export of alkaline phosphatase, phospholipase C, elastase and exotoxin A (Strom et al., 1991). These extracellular products are virulence factors critical for bacterial infection. On the other hand, as cleavage of signal peptide by PilD is necessary for the pilus assembly (Strom et al., 1993), PilD mutations result in type IV-specific defections (Fig. 2). Type IV system in *P. aeruginosa* influences many physiological processes, including adherence to living surfaces, twitching motility, modulation biofilm biosynthesis, DNA uptake and exchange and exoproteins secretion (Giltner et al., 2012). It can expect that the growth and development of PilD mutant subpopulation in the whole community would largely determine the population virulence.

**Figure 2.**
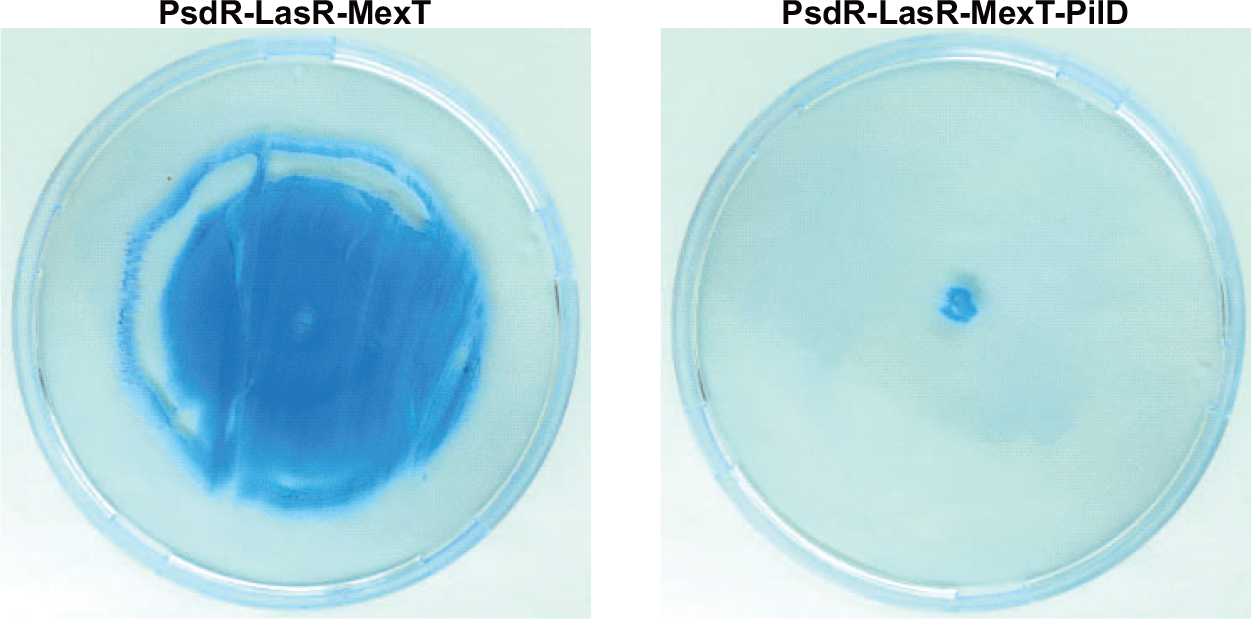
Twitching motilities of PilD mutants. Twitching motilities of PsdR-LasR-MexT and PsdR-LasR-MexT-PilD mutants were assayed. Thin agar plates (1.0%) were stab inoculated with a toothpick to the bottom of the plate and incubated for 24 h at 37 ° C. The twitching zone was visualized by Coomassie Blue staining.

## Materials and methods

### Bacterial strains and growth

The PAO1 strain and the mutant derivatives were grown in Luria Bertani (LB) broth containing 10 mg/ml tryptone, 5 mg/ml yeast extract, 10 mg/ml NaCl at 37 °C or photosynthetic medium (PM) (Kim and Harwood, 1991) supplemented with 1% sodium caseinate (Sigma Aldrich, USA) as the sole carbon source at 37 °C. Unless otherwise specified, *P. aeruginosa* strains were cultured in 14-mm FALCON tubes (Corning, USA) containing 3 mL medium, with shaking (225 RPM) at 37 °C. Escherichia coli was grown in LB broth at 37 °C.

### Evolution experiments

Two independent colonies of PsdR-LasR-MexT mutants were separately inoculated into 3 mL LB-Mops broth for overnight culture. Cell evolution was initiated by transferring 50 μl bacteria grown in LB-Mops into 3 mL PM medium in 14-mm FALCON tubes (Corning, USA). Passage was performed every 5 days by exchanging 50 μl bacteria into 3 mL fresh PM medium. To identify protease-positive variants, evolved colonies were isolated on LB agar and spotted onto the skim milk plates at 37 °C for 18h to monitor the protease-catalyzed zones.

### Construction of PilD mutant

Full-length of *pilD* gene knocking out was based on the homologous recombination exchange approach as described previously. Briefly, about 500 ~ 1000 bp of DNA flanking the targeted full length of gene of interest were PCR-amplified and cloned into pEXG2 vector (Gentamycin resistance, Gm) with the Vazyme ClonExpress II One Step Cloning kit (Vazyme Biotech, Nanjing, China), generating pEXG-flanking constructs. The pEXG-flanking construct was mobilized into *P. aeruginosa* strain by triparental mating with the help of *E. coli* PRK2013 strain (Kanamycin resistance, Km). Deletion mutants were first selected on Pseudomonas Isolation agar (PIA) containing 100 μg/ul Gm and further selected on LB agar containing 10% sucrose. All mutants were confirmed by PCR amplification and subsequent DNA Sanger sequencing.

### Skim milk plate assay

Total proteolytic activities of P. aeruginosa strains were evaluated through skim milk assay, in which the tested strains form a zone of clearing on skim milk agar plate. Individual colonies were spotted on the skim milk agar plates (25% LB, 4% skim milk, 1.5% agar). The protease-catalyzed zones were photographed after incubation at 37 °C for 16h.

### Twitching motility

Surface-associated twitching motility was assessed via agar stab inoculation in 1% LB agar plate. Colonies were transfer from solid LB agar plate and stab inoculated in the 1% LB agar plate. Twitching motility zones (occurring at the interface of agar/petri plate) were allowed to develop 24 h at 37 °C, after which the agar was removed. The twitching zones were then visualized by staining of 0.06% Coomassie Brilliant Blue and 10% Acetic Acid for 10 min and washing with distilled water briefly. Skim milk assay

## Figure legends

**Supplementary Tables**

**Table S1. Identification of mutations in *pilD*.**

