## Supplementary Table S1 for "PilD mutant is a new cheater in *Pseudomonas aeruginosa* quorum-sensing evolution"

### Table S1. Identification of mutations in *pilD*.

| Evolved colony | Nucleotide change | Encoding | Targeted gene | Product |
|---|---|---|---|---|
| PsdR-LasR-MexT#1 | deletion 5073010-5073025 (16 bp) | CDS | *pilD* | prepilin peptidase |
| PsdR-LasR-MexT#2 | deletion 5073323 (1 bp) | CDS | *pilD* | prepilin peptidase |

Whole-genome re-sequencing (WGS) was performed with the evolved protease-negative colonies from the PsdR-LasR-MexT population. Mutations were identified relative to the reference *P. aeruginosa* PAO1 genome (NC_002516.2).
